# Time-resolved mapping of genetic interactions to model rewiring of signaling pathways

**DOI:** 10.1101/384800

**Authors:** Florian Heigwer, Christian Scheeder, Thilo Miersch, Barbara Schmitt, Claudia Blass, Mischan Vali Pour-Jamnani, Michael Boutros

## Abstract

Context-dependent changes in genetic vulnerabilities are important to understand the wiring of cellular pathways and variations in different environmental conditions. However, methodological frameworks to investigate the plasticity of genetic networks over time or in response to external stresses are lacking. To analyze the plasticity of genetic interactions, we performed an arrayed combinatorial RNAi screen in *Drosophila* cells at multiple time points and after pharmacological inhibition of Ras signaling activity. Using an image-based morphology assay to capture a broad range of phenotypes, we assessed the effect of 12768 pairwise RNAi perturbations in six different conditions. We found that genetic interactions form in different trajectories and developed an algorithm, termed MODIFI, to analyze how genetic interactions rewire over time. Using this framework, we identified more statistically significant interactions compared to endpoints assays and further observed several examples of context-dependent crosstalk between signaling pathways such as an interaction between Ras and Rel which is dependent on MEK activity.

## INTRODUCTION

Gene-gene interactions, the epistatic influences of one gene’s effect on the function of another gene, have widespread effects on cellular and organismal phenotypes – ranging from fitness defects in unicellular organisms to interactions between germline and somatic variants in cancer (Baryshnikova et al., 2013; Billmann and Boutros, 2017; Boone et al., 2007; Burgess, 2016; Carter et al., 2017; Ideker and Krogan, 2012; Mani et al., 2008; Phillips, 2008; Taylor and Ehrenreich, 2015). In past studies, statistical genetic interactions (also simply referred to as genetic interactions) have been defined as an unexpected phenotypic outcome observed upon simultaneous perturbations (or knock-outs) of two genes that cannot be explained from the genes’ individual effects (Beltrao et al., 2010; Fisher, 1930; Mani et al., 2008).

Genetic interactions have been discovered using pairwise perturbations of genes, a strategy which has been experimentally used at large scale in yeast (Collins et al., 2007; Costanzo et al., 2010; Fiedler et al., 2009; Tong et al., 2001a), *C. elegans* (Lehner et al., 2006), *Drosophila* (Fischer et al., 2015; Horn et al., 2011), *E. coli* (Babu et al., 2011) and human cells (Kampmann et al., 2013; Laufer et al., 2013; Roguev et al., 2013; Shen et al., 2017). To create genetic interaction maps, these studies systematically identified positive (e.g. better fitness than expected) or negative genetic interactions which can then be used to generate ‘genetic interaction profiles’ for each gene. Several studies have shown that profile similarities are a powerful predictor for gene function and they have been used to create maps of cellular processes at a genome-wide scale (Costanzo et al., 2010, 2016; Fischer et al., 2015; Pan et al., 2018; Rauscher et al., 2018; Tsherniak et al., 2017; Wang et al., 2017; Yu et al., 2016).

In addition to univariate phenotypes, such as fitness and growth phenotypes of cells or organisms, genetic interactions can be measured for a broader spectrum of phenotypes by microscopy and image-analysis (Horn et al., 2011; Laufer et al., 2013; Roguev et al., 2013). Importantly, multi-variate phenotypes further opened the possibility to predict the epistatic relationship of components in genetic networks by deriving the direction of specific genetic interactions (Fischer et al., 2015).

Most studies of genetic interactions have been performed under ‘static’ environmental conditions. In contrast, several studies have analyzed the impact of environmental changes on genetic interaction networks. This way, they investigated how interactions differ between steady states in different environmental conditions (Bandyopadhyay et al., 2010; Billmann et al., 2017; Díaz-Mejía et al., 2018; Guénolé et al., 2013; Martin et al., 2015; St Onge et al., 2007; Wong et al., 2015). For example, Bandyopadhyay *et al.* (2010) defined static, positive and negative differential interactions that vary under changing environmental conditions. And, Billmann *et al.* (2017) used extrinsic and intrinsic changes of Wnt signaling in cultured *Drosophila* cells to map differential genetic interactions using a pathway-directed phenotypic readout. These studies showed that upon changes in the environment widespread changes in genetic interactions occur.

Upon treatment, e.g. with small molecules, genetic interactions change over time due to time-dependent depletion of components or other changes in the underlying composition of its molecular constituents. However, to date little is known about how genetic interaction networks ‘rewire’ over time and models for their analysis as well as proof-of-principle data sets are missing. In this study, we devised an experimental and analytical approach to gain insights into higher order (e.g. gene-gene-drug) differential interactions. To analyze how they manifest over time we used an image-based, multi-variate phenotypic readout. By combining gene combinatorial RNAi with a MEK inhibitor or control treatment, we measured higher order chemo-genetic interactions in *Drosophila* S2 cells.

In this study, we first performed image-based genome-wide RNAi screens to identify a gene set that modulates the phenotypic profiles upon MEK inhibition. To construct the differential genetic interaction network, we then created a double-perturbation matrix and measured the effect of 12768 gene-gene perturbations under time and treatment differential conditions. These perturbations span a previously defined set of 168 x 76 genes and were characterized by 16 phenotypic features. Notably, we assessed how each differential interaction changes over time and used this information to construct maps of treatment-responsive biological modules. Differential interactions mapped the plasticity of Ras signaling and crosstalk to other signaling pathways, such as *Rel* and Stat signaling. Our analyses should help to better understand the principles of interaction changes in higher order combinations of genetic perturbations.

## RESULTS

Ras signaling is an important oncogenic pathway and its upstream members Ras and EGFR family proteins are frequently mutated in cancer (Rodriguez-Viciana et al., 2005). MEK1/2 (the human ortholog of *Dsor1*) acts downstream of Ras and phosphorylates ERK1/2 (the human ortholog of *rl*) which itself phosphorylates many other proteins (e.g. ETS-family transcription factors) mediating mainly proliferative signals (Friedman et al., 2011). The topology of the Ras signaling pathway and its key components are largely conserved between human and *Drosophila* (Kolch, 2005; Perrimon, 1994; Wassarman et al., 1995). In *Drosophila*, the Ras-pathway has been implicated in the growth of wing imaginal discs, differentiation of photoreceptors and hemocyte proliferation and is one of the main mechanisms controlling cell proliferation (Asha et al., 2003; D’Neill and Rubin; Prober and Edgar, 2000; Wassarman et al., 1995). Willoughby *et al.* (2013) previously compared the effect of multiple MEK-small molecule inhibitors *in vivo* and S2 cell culture and showed that all but one inhibitor significantly reduced the levels of phosphorylated *rl*.

To recover a broad spectrum of cellular phenotypes upon MEK-inhibition, we used an adapted cell morphology assay and automated image analysis in *Drosophila* cells (Breinig et al., 2015; Fischer et al., 2015; Horn et al., 2011). In this assay, we perturbed cells by small molecule treatment or genetic perturbagens, before we stopped the cell-based assay by fixation and staining for DNA (visualizing the nucleus), actin (visualizing cell adhesion and cytoskeleton organization) and α-tubulin (visualizing cell cycle dynamics). Automated high-throughput microscopy combined with a real-time image analysis framework then scored phenotypes on a single-cell level. The resulting multi-variate phenotypic feature vector describes the quantitative phenotype resulting from the perturbation (Figure 1A, see Methods), also referred to as a phenoprint.

**Figure 1.**
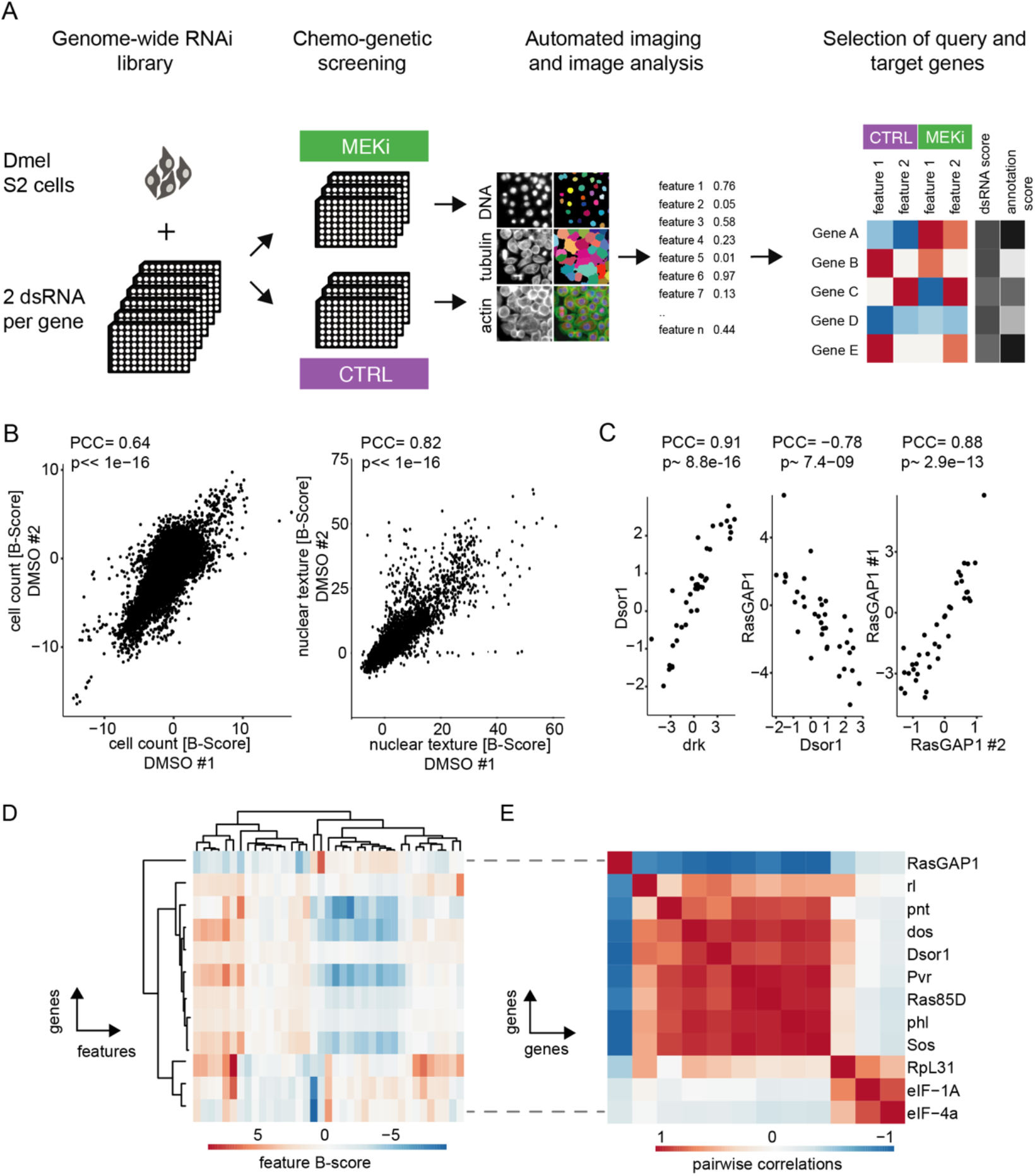
Measuring chemo-genetic interactions by high throughput imaging and RNAi. (A) Schematic representation of the workflow. A genome-wide RNAi library covering 13617 unique *Drosophila* genes with two dsRNA constructs was screened in the presence of DMSO (control) or MEKi (PD-0325901) at 1.5 nM. Fixed cells were stained for DNA, actin and α-tubulin and imaged at 20x magnification at a resolution of 2048×2048 pixels. Phenotypes were analyzed by automated image analysis of cell morphology on a single-cell level using an R/EBImage pipeline. Genes were rank ordered for co-RNAi screening by dsRNA score (quality and differential phenotype) and gene annotations (unknown genes favored). (B) Well-level correlation of biological replicates. Shown are B-score normalized cell count data (left, PCC=0.64) and nuclear texture scores (right, PCC=0.82) exemplarily for the control condition. One outlier was flagged and not plotted for nuclear texture. (C) Comparison of induced phenotypes by correlation of feature vectors (Pearson correlation coefficient [PCC] and asymptotic p-value). Knockdown of Dsor1 and drk, two known positive regulators of Ras signaling resulted in high (PCC=0.91) positive correlation. Knockdown of RasGAP1 (neg. regulator of Ras) and Dsor1 induced inverse correlating feature vectors (PCC=−0.78). dsRNA reagents directed against the same gene show a high level of similarity (RasGAP1 design #1 vs. #2, PCC=0.88) (D) Feature vectors for selected genes regulating Ras signaling and protein translation. Hierarchical clustering groups genes into regulatory units, marked by similar feature profiles. Shown are average B-scores across replicates of 16 representative features. (E) Hierarchical clustered matrix of pairwise correlations. Pairwise PCC were calculated for all example genes in (D). Clustering genes by their pairwise correlation sorted them into groups of functionally related genes. All data show MEK inhibited condition, if not noted otherwise.

We confirmed the suitability of this multi-variate cell-based assay to score compound induced phenotypes without the need to measure its direct biochemical effect (such as rl phosphorylation). We further used the assay to determine the ED50 of the MEK-inhibitor PD-0325901 on Dmel2 cells (**Supplemental Figure S1**). These experiments demonstrated that Dmel2 cells show a sustained phenotypic response towards PD-0325901. A high correlation (Pearson’s correlation coefficient [PCC]=0.81) between small molecule and RNAi perturbation of MEK indicates high compound specificity (**Supplemental Figure S1**).

### A chemo-genetic screen identifies MEK inhibitor-sensitive genes

As combinatorial gene perturbation screens scale poorly with the number of genes, we first sought genes which phenotypes change in a drug-dependent manner. Previous studies have found that gene-gene interactions are enriched for candidate genes that display a phenotype itself (Deshpande et al., 2017; Koch et al., 2017). Hence, the identification of genes showing a phenotype as a single knockdown will in turn likely enrich combinatorial screens for genes that form higher order interactions. To map gene-gene interactions sensitive to MEK inhibition, we performed multiple genome-wide RNAi screens under different environmental conditions (Figure 1A).

We used the HD3 RNAi library, which targets 13617 genes (unique FlyBase IDs) in the *Drosophila* genome with two independent double-stranded (ds) RNAs (see Methods). We reverse transfected Dmel2 cells with this genome-wide library in 384-well plates and treated them after 24 h with the PD-0325901 or DMSO as a control. After 3 additional days, we fixed, stained and imaged the cells. We collected data from 360 384-well micro well plates with four fields of view per well in three fluorescence channels, generating a dataset of 1.6 Mio. images. Screens were conducted in biological replicates. We then used automated image analysis to segment ~10000 cells per well and calculated features for individual cells’ nuclear and cytoplasm shape, texture, intensity profiles (see Methods, Breinig et al., 2015; Fischer et al., 2015a; Laufer et al., 2013a). 38 phenotypic features that displayed a high correlation between biological replicates were used for further analysis (**Supplementary Table S1**, replicate correlation PCC>0.5).

To assess the quality of our profiling assay we first analyzed overall reproducibility. For example, the screens showed a correlation between biological replicates of PCC=0.64 and PCC=0.82 for cell count and nuclear texture, respectively (Figure 1B). Phenotype feature vectors also reliably separated control RNAi treatments (*RasGap1* [*RASA3*] vs. *drk* [*GRB2*], multi-variate Z’=0.695, **Supplemental Figure S2**). In addition, the multi-variate Z’ factor is significantly higher than univariate Z’ using cell count only (0.695 > 0.295, p<0.01, Zhang et al., 1999). We also found that the phenotypes produced by knockdown of known Ras pathway components *Dsor1* (*MEK1/2*) and *drk* showed a high correlation (PCC=0.91, Figure 1C). Accordingly, knockdown of genes with antagonizing biological function like the negative regulator of Ras signaling *RasGAP1* (*RASAL3*) and *Dsor1* (Feldmann et al., 1999) resulted in phenotype vectors that inversely correlate (PCC=−0.78). dsRNA targeting the same gene were also highly reproducibly producing similar phenotypic vectors (e.g. PCC_RasGAP1_=0.88).

Next, we calculated average phenotype vectors for the core Ras signaling cascade, its negative regulator RasGAP1 and genes involved in the translation machinery, as a control (Figure 1D). Hierarchical clustering recapitulated known functional relationships of Ras pathway components, whereas translational regulators show clearly distinct phenotype vectors (Figure 1E). These experiments demonstrated that the morphology assay is able to capture functional meaningful phenotype vectors for MEK inhibition, and robustly distinguishes control perturbations. It also groups functionally related or antagonistic genes into clusters of phenotypic similarity or dissimilarity, respectively.

In order to generate a focused library to profile gene-gene interactions, we selected a set of 168 genes from these screens that showed: (i) high reproducibility between biological replicates, (ii) high correlation between sequence independent dsRNA reagents, (iii) measurable effects that deviate from the negative controls and (iv) differential phenotypes upon *Dsor1* inhibition (**see Supplemental Table S2**). We also prioritized genes that were largely uncharacterized (see Methods). The resulting gene list for gene-gene interaction screening covers 168 target genes with highly reproducible phenoprints between biological replicates and dsRNA reagents (PCC>0.5) that are expressed in Dmel2 cells (log_2_ (RPKM+0.001) > 0) and show phenotypic differences between control and treated conditions. These genes also cover a number of signaling pathways including Ras signaling, innate immunity, Wnt signaling, mRNA splicing, protein translation, cell cycle regulation, Jak/STAT signaling and Tor signaling (**see Supplemental Table S3**). The query gene set, a subset of the target genes, contained 76 well described genes to aid biological interpretability.

### A time resolved co-RNAi screen to capture differential genetic interactions

Previous studies defined positive differential, negative differential and stable interactions between two genes associated to changes in environmental conditions such as DNA-damage (Figure 2A, Bandyopadhyay et al., 2010; St Onge et al., 2007). Positive differential interactions are newly forming under stress conditions and mark resistance or other mechanisms counteracting the noxious stimulus (e.g. drug treatment). Negative differential interactions, on the contrary, mark connections that are required for homeostasis under normal, unperturbed conditions but are either obsolete or harmful under stress conditions. Within these studies, the wiring diagrams of genetic interaction networks were studied at steady state conditions between two end-points. The information gained from observations of isolated gene-gene-drug interactions thus missed dynamic responses of differential interactions (Bandyopadhyay et al., 2010; Ideker and Krogan, 2012; Mani et al., 2008; Martin et al., 2015).

**Figure 2.**
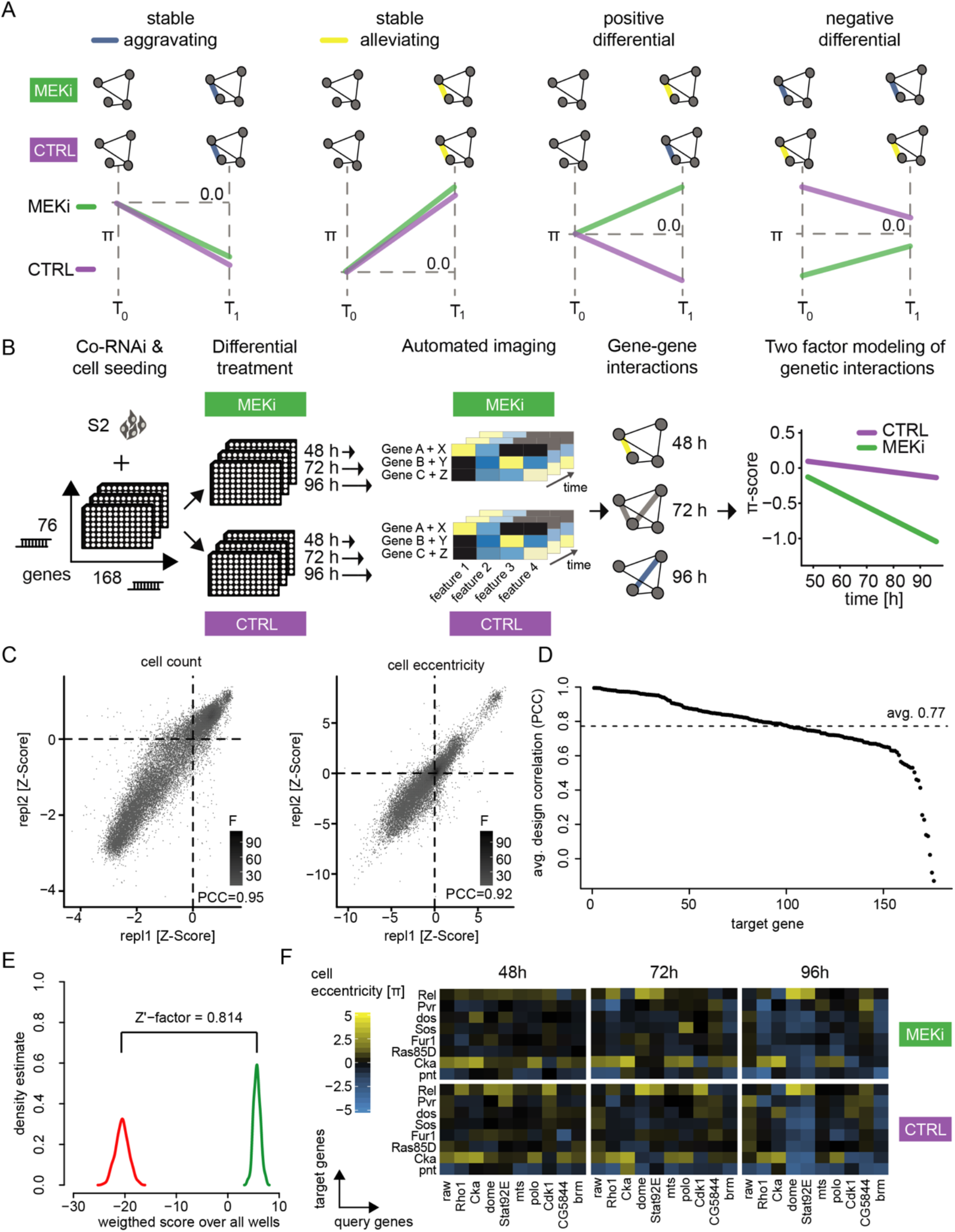
Genetic interactions screening using RNAi yields highly reproducible results. (A) Types of differential genetic interactions shown between T_0_ and T_1_. Alleviating interactions shown in yellow and aggravating interactions shown in blue: treatment invariant but time dependent interactions (stable) and interactions which are treatment differential depending on the time point (differential). (B) Representation of the combinatorial RNAi (co-RNAi) screening setup. 168 ‘target’ and 76 ‘query’ genes were combined to all pairwise combinations and arranged accordingly in 384-well plates. S2 cells were reverse transfected with pre-spotted dsRNAs and incubated for 24 h. Cells were treated either with small molecule (MEKi [PD-0325901], 1.5 nM) or DMSO (solvent control, 0.5 % DMSO) and incubated for additional 48 h, 72 h or 96 h. The assay was stopped by fixation and staining of cells. Phenotypes were measured using automated microscopy and image analysis. Genetic interactions (π-scores) were called for 16 non-redundant phenotypic features from the combinatorial treatments, separately for each treatment and timepoint. MODIFI was applied to identify significant differential genetic interactions. The model is defined as π_[A,B,time,treatment]_ ~ σ_[A,B]_ * *time* + δ_[A,B]_ * *treatment* + ε_[A,B]_ with π being the measured interaction for a pair of genes A and B at a given time and treatment. (C) Quality control of the co-RNAi screen. Scatter plots showing all replicate measurements of normalized and processed phenotypic feature data for cell count (PCC=0.95) and cell eccentricity (PCC=0.92). Plates for which positive (RasGAP1) and negative (Diap1) controls could not be distinguished (Z’-factor < 0.3) were excluded. Data was normalized for plate and batch effects using median normalization and scaled and centered using *Z*-score analysis. (D) RNAi reagents ranked by dsRNA design correlation. dsRNAs were paired by gene to calculate pairwise correlations along 16 selected features. All dsRNAs together showed an average design correlation of PCC=0.77. Figure C and D show data 96 h after MEK inhibitor treatment representatively. (E) Assay performance assessed by statistical effect size. The multi-variate Z’-factors between the positive control knockdown of RasGAP1 and a negative control knockdown of Pvr (Zhang et al., 1999) were calculated for the selected 16 phenotypic features for the MEKi screen after 96 hours. (F) Example of genetic interactions observed over time and treatment. Interaction data for the inhibitor treated and control condition are shown for eight selected target genes (y-axis) and 10 query genes (x-axis). Genetic interactions shown were calculated for the cell eccentricity feature.

Next, we aimed to quantitatively analyze differential genetic interactions in a time dependent manner and set up an experimental design based on co-RNAi treatment and high-throughput microscopy (Figure 2B). A combinatorial gene-gene matrix covering 168 target genes and 76 query genes was used to measure 12768 genetic interactions under the different conditions. The library was screened under MEK (Dsor1) inhibitor and control conditions at 48, 72 and 96 hours after compound addition. The screen was performed using two sequence-independent dsRNA design replicates and in two biological replicates for each condition. 4.4 Mio. fluorescent images were captured, and 155 image features measure the perturbation effects for every single cell (see **Supplementary Methods**). Following automated image analysis, we transformed the phenotypic features using the generalized logarithm, normalized, centered and scaled them (see Methods). Plates failing technical quality control (Z’-factor between *RasGAP1* RNAi and Diap1 RNAi < 0.3 and biological correlation < 0.6 PCC for cell number) were masked in further analysis. Overall, < 3% of all plates were excluded according to these criteria. Most of the 155 features showed a high reproducibility (80 % having a PCC greater than 0.6, MEKi condition) (**Supplementary Figure S3A**). The two features cell count (relative cellular fitness) and actin eccentricity (morphology of cells) were among the features with the highest replicate correlation (Figure 2C) and are highlighted as exemplary features in some of the following visualizations. All features that failed to meet a replicate correlation of PCC > 0.6 were removed, leaving 114 features for further analyses (**Supplementary Figure S3A**). In addition, 90 % of sequence-independent dsRNA pairs correlate with a PCC > 0.6 with an average correlation of PCC = 0.77 (Figure 2D).

Since many of the remaining 114 features provide redundant information (**Supplementary Figure S3B**), overlap was reduced by first clustering all features according to the pairwise PCC of the genetic interactions. Second, we fixed the first feature (cell number) and removed all remaining features that correlated with PCC > 0.7. Third, we selected the next most reproducible and biologically interpretable feature and removed all highly correlated features; this scheme was iterated until all features were passed. The remaining 16 features (see **Supplementary Table S4**) were selected for further analysis. As a confirmation, we verified that cell number and actin eccentricity show a weak correlation (PCC = 0.48) and thus provide independent information (**Supplementary Figure S3C**). An unbiased information gain analysis by stability selection, as carried out in an earlier study (Fischer et al., 2015), validated this approach showing that each of the chosen features also delivers independent but reproducible information (**Supplementary Figure S3D**). As they enrich biologically interpretable and reproducibly measurable features we, however, kept the features selected by correlation-based analyses. An analysis of the multi-variate Z’-factors between RasGAP1, a negative regulator of Ras signaling and Pvr, a positive regulator of Ras signaling (Zhang et al., 1999) revealed a multi-variate Z’ of 0.814, indicating high assay quality (Figure 2E).

Following quality control, we calculated genetic interaction scores (π-scores) for each feature under each condition using a multiplicative model as described previously by Horn *et al.* (Horn et al., 2011, see Methods). **Supplementary Figure S4** exemplifies this approach for the interaction between two example genes. Overall, we analyzed of 1.3 million gene-gene interactions in two conditions, three time points and 16 cellular features. 72922 interactions showed a significant deviation from the expected combinatorial phenotype. Only 9090 (12%) genetic interactions are measured significantly (moderated t-test [limma], FDR<0.1) for the cell number phenotype.

Next, we systematically analyzed whether: (i) π-score analysis recapitulates earlier studies using a morphology readout in *Drosophila*, (ii) π-scores were reproducible between biological replicates, (iii) the interaction profile changed considerably when target and query genes switch roles and (iv) interaction profiles were independent for different features. To this end we compared gene-gene interactions that overlapped between this and previous studies of genetic interactions in *Drosophila* S2 cell culture (**Supplementary Figure 5**). We found significant agreement between π-scores measured in various features in the different studies (FDR<<0.1, for linear dependence between π-scores measured in different studies). We found, for example, that the DNA texture feature we used could also explain the phospho-histone H3 staining used in Fischer *et al.* (2015).

Next, we confirmed a high correlation of interactions between biological replicates, as illustrated on the phenotypic features ‘DNA eccentricity’ and ‘cell number’ (**Supplementary Figure S6A, A’**). As the combinatorial matrix contained all query genes also in the target gene set, we tested whether interaction phenotypes were in accordance regardless of the assignment of target and query. In theory, all interactions should be symmetric, and it should not matter which gene was assigned as target and which as query. However, in practice target and query RNAi reagents were added independently during the experiment which could skew symmetry. Our analysis demonstrated that both combinatorial conditions highly correlate (**Supplementary Figure 6B, B’**, PCC= 0.76 for cell number; PCC=0.75 for actin eccentricity). We furthermore confirmed that different features provide independent information about genetic interactions as indicated by low correlation (PCC=−0.21 and 0.04, **Supplementary Figure 6C, C’**).

### Robust linear modelling of differential genetic interactions across multiple features

Figure 2F shows an excerpt of the genetic interaction matrices obtained for each treatment and time condition. We found that our analyses recapitulated known genetic interactions. For example Ras signaling components showed negative interactions with Jak/STAT pathway (e.g. *Pvr, dos* and *Sos* show negative genetic interactions with *dome* and *Stat92E* (*STAT5B*), Baeg et al., 2005; Li et al., 2003; Xu et al., 2011). The observed interactions become stronger over the three time points measured, and interactions such as a negative interaction between Ras signaling components and Rho1 are stronger upon MEK inhibitor treatment.

Next, we sought a suitable statistical framework to score significant differential interactions. Previous studies employed different statistical tests that score the significance of interaction differences between end-point measurements (*B-Score, dS-Score*, limma based moderated t-test, Bandyopadhyay et al., 2010; Bean and Ideker, 2012; Billmann et al., 2017; Guénolé et al., 2013). In a pooled genetic interaction screen in human cells, Shen *et al.* used the time dependence of fitness defects to improve statistical power (Shen et al., 2017). Thus, we tested whether we can also leverage a time and treatment dependent model (Figure 3A) to identify differential genetic interactions more sensitively than time independent statistical models (Figure 3B).

**Figure 3.**
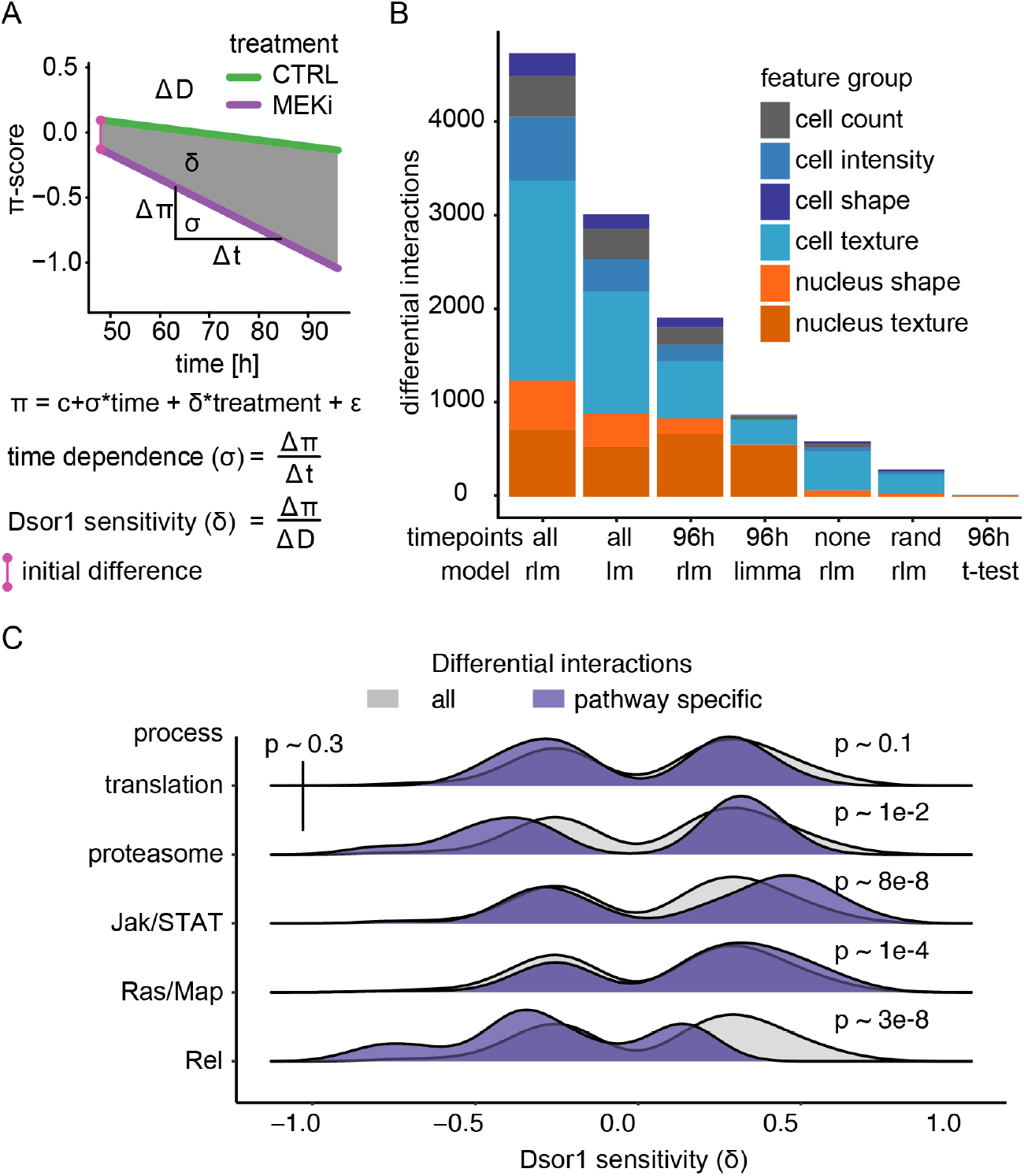
Robust linear models describe the continuity of genetic interaction rewiring. (A) Derived measures from the interaction model: the speed (σ) of interaction development, amplitude (*δ*) of response to MEKi and average initial interaction difference. (B) Differential interaction counts depending on sequential data and the statistical model applied. Significant (FDR < 0.1) differential interactions were counted when analyzed using conventional linear modeling, robust linear modeling, moderate t-test as of R/limma or student’s t-test. End-point, sequential and randomized data were compared. The analysis was carried out for all features and accumulated counts are shown, “none” means that all time points were treated as replicates of the same measurement, “rand” means that measurements were assigned to random time points and 96 h denotes the data treated as end-point measurements of the last time point. All models tested the NULL hypothesis that there is no difference between treatments. A two sided welch t-test was used. The robust linear model (rlm) coefficient’s significance was estimated using robust f-tests. The linear model (lm) coefficient’s significance was tested by two-way ANOVA. (C) Measurement of sensitivity towards MEKi. The sensitivity to Dsor1 inhibition was assessed by comparing *δ* between various biological processes. Significance was tested by a two-sided Kolmogorov-Smirnov test of the sample against all measured interactions. Resulting p-values are indicated.

To this end, we tested whether the development of genetic interactions over time and between different conditions could be quantitatively described by a multi factorial linear model. This would provide the possibility to (i) quantify the time dependence of an interaction and (ii) to measure the phenotypic difference between treatment conditions with high confidence. For every gene-gene combination [i, j], we used a two-factor robust linear model, which we termed model of differential interactions (MODIFI), to estimate the predictive strength and influence of time and differential compound treatment on the π-score.

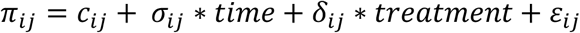

Therein, the coefficient σ_ij_ models the time dependence, δ_*ij*_ models the quantitative offset between treatments, c aids modeling without interpretation as the intercept and the residual ε_ij_, estimates the error of fit for each combination of the target gene i and the query gene j. σ and δ are thus parameter estimates that uniquely describe the behavior of each gene-gene interaction.

We found that the robust linear model of serial measurements (MODIFI) identifies the most differential interactions (4723 in total, 2.31 % of all possible interactions, FDR<0.1). When using only end-point measurements, the robust statistic (rlm) is more sensitive than the moderated t-test (limma) used in previous studies to score differential genetic interactions (Billmann et al., 2017; Fischer et al., 2015; Laufer et al., 2013) and the two-tailed t-tests of each interaction between conditions (1907 vs 874 vs 21 interactions, respectively; Bandyopadhyay et al., 2010; Guénolé et al., 2013). We further found that MODIFI increased statistical power by identifying 147 % more differential interactions across all features when compared to the best end-point measurements (4723 vs 1907; 96h/rlm). We conclude that by employing robust statistics MODIFI outperforms conventional models and more accurately estimates the parameters δ and σ.

Next, we inferred to what extend gene-gene interactions changed due to the MEK inhibitor treatment. This parameter (δ) serves as a surrogate for the integrated area between the trajectories of the two treatments. If δ is close to zero, only little changes occur upon treatment and *vice versa*. We found that differential interactions were equally likely to be positive or negative differential over all analyzed genes (Figure 3C, grey distribution). Of note, especially differential interactions of *Rel (NF-kB, downstream effector* of the *Drosophila* Imd signaling pathway, Myllymäki et al., 2014), or Ras/Map and Jak/STAT related genes enriched as either negative (π-score declines because of MEK inhibition) or positive (π-score rises because of MEK inhibition), respectively. This implies that pathways, which are positively regulated by MEK, tend to form interactions that are lifted (less aggravating) under MEK inhibition. Interactions formed by *Rel* seem the be negatively enhanced by MEK inhibition. We further found no or little significant difference between housekeeping modules (proteasome, translation machinery) and all measurements (p>0.1, two-sided KS-test). Taken together, these data suggest that components of the same pathway share differential interaction sensitivity and directionality in response to Ras pathway inhibition.

### Differential genetic interactions enrich in stress responsive genes and pathways

Assessing all interactions for which MODIFI produced a statistically significant fit (FDR<0.1), we identified four main types of time and treatment dependent interactions that we expected would be recovered by this experiment (also compare Figure 2A). We observed alleviating stable interactions when the π-score raised over time and was indifferent between treatments (Figure 4A). Positive stable interactions often involve core essential genes whose influence on the phenotype (e.g. cell count) is not altered by MEK inhibition. This is, for example, the case for *mts* knockdown (PP2CA, lethal by itself; Snaith et al., 1996) where the simultaneous loss of the proteasomal subunit *Prosbeta4* (PSMB2, Wójcik and DeMartino, 2002) dominates the combinatorial phenotype that cannot be worsened regardless of the treatment. A positive interaction that strengthens over time is measured (Figure 4A). Accordingly, we termed it an aggravating stable interaction when the π-score declined over time and its trajectories were indifferent between treatments (Figure 4B).

**Figure 4.**
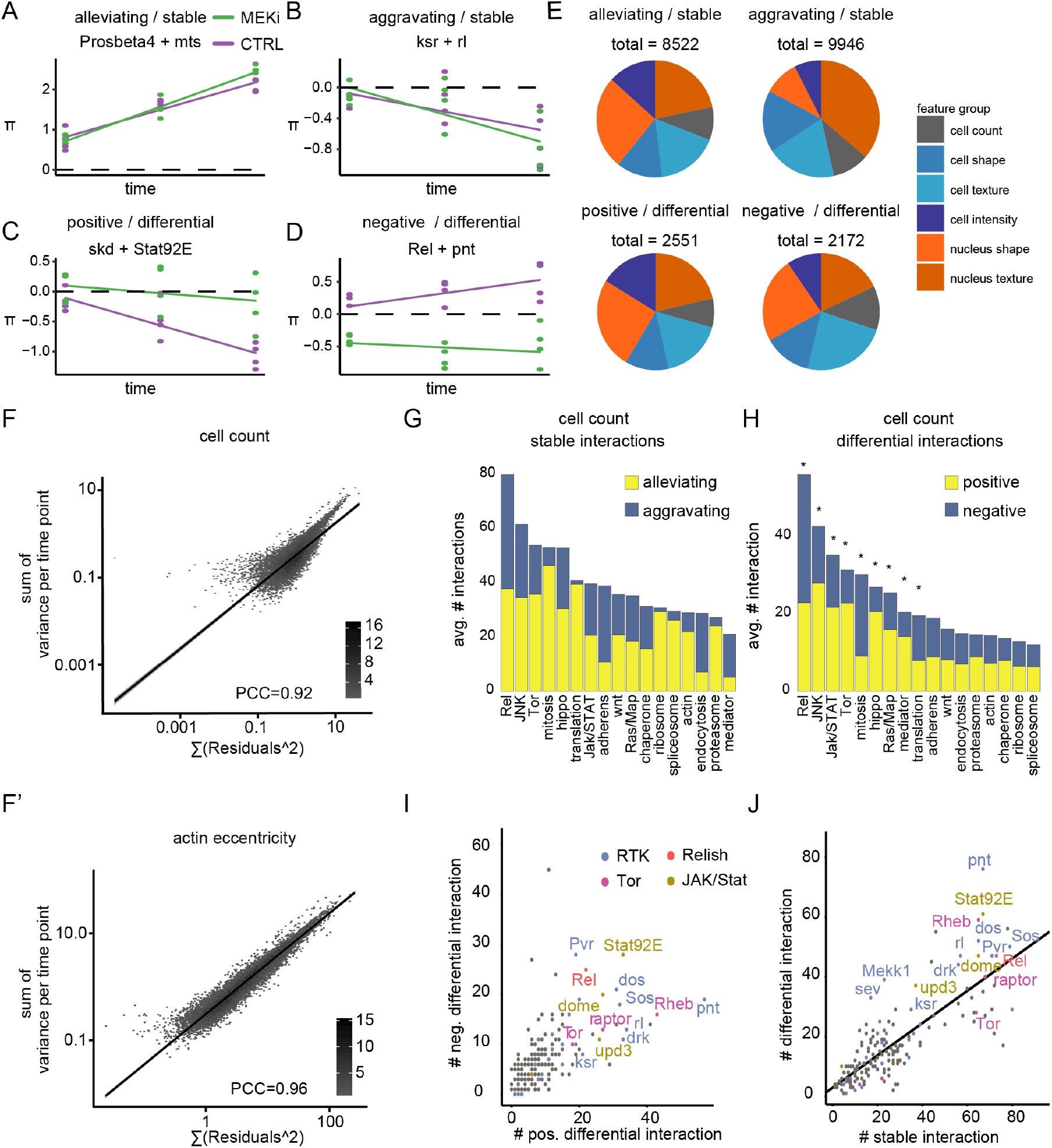
Genetic interactions rewire over time. (A-D) Examples of time and treatment dependent genetic interactions: (A) alleviating-stable interaction of *Prosbeta4* (proteasome) and *mts* (cytoskeleton), treatment invariant and increasing over time, (B) aggravating-stable interaction of *ksr* and *rl* (both Ras signaling), treatment invariant and decreasing over time, (C) positive differential interaction of *skd* (mediator complex) and *Stat92E* (STAT receptor), lifted from synthetic lethal to non-interacting by treatment, (D) negative differential interaction of *Rel* (innate immunity) and *pnt* (Ras signaling), π-scores decreased by the treatment. (cell count, FDR < 0.1, robust f.test + Benjamini Hochberg). (E) Interaction counts after MODIFI. Interactions (FDR < 0.1) are counted for 16 features, grouped into cell count, shape, texture and intensity within cell and nucleus. (F) Robust linear model credibility. Residuals are compared to replicate variance for each time point, exemplarily shown for cell count (F, PCC=0.92) and cell eccentricity (F’, PCC=0.96). Trendline indicates good agreement of residuals and variance. (G-H) Distribution of aggravating/alleviating stable (G) and positive/negative differential (H) interactions among molecular pathways. Binomial testing estimated if counts were expected by chance (*~FDR<0.1). (I) Gene-level interaction counts. Counts of significant, unique negative differential interactions compared to counts of positive differential interactions. Dots are colored by functional groups. Pathways with the most differential interactions (Tor, Ras, Rel and Jak/STAT signaling) are highlighted. (J) Counts of stable interactions are plotted against treatment differential interaction counts. A trendline indicates a general linear dependency between stable and differential interaction counts. (G-J) Count data are based on cell number feature, significant (FDR < 0.3), MODIFI modelled interactions.

Aggravating stable interactions on cell count often include signaling transducers where the loss of one only has a mild phenotype while the double perturbation disturbed homeostasis to an extend the cells cannot buffer and synthetic lethality is observed. For example, *ksr* (*KSR1*) and *rl* (*ERK1/2*), two core members of the Ras signaling cascade downstream of Dsor1 (Morrison, 2001; Wassarman et al., 1995), interact significantly (p~0.0017). This interaction is stable upon MEK inhibition and thus appears independent of phospho-rl levels (Figure 4B, **Supplemental Figure 1**). This indicates that *ksr* and *rl* form a synthetic sick interaction that influences cell viability independent on the Ras signaling phosphorylation cascade.

We defined interactions as differential when trajectories differed significantly between treatments (FDR<0.1). If the π-score is lower under control then under treatment conditions, we termed it a positive differential interaction (MEK inhibition lifts the phenotype, Figure 4C) and negative differential interaction (MEK inhibition dampens the interaction, Figure 4D) in the opposite situation. For instance, *skd* (MED13, an integral component of the mediator complex; Janody et al., 2003) showed a positive differential interaction with *Stat92E* (*Drosophila* ortholog of human STAT receptor; Bina and Zeidler, 2013) (Figure 4C). Under control conditions *skd* knockdown aggravated the fitness loss induced by *Stat92E* knockdown. This aggravation was attenuated under MEK inhibition. Our data suggest that a synthetic lethal relationship connects both genes when they are otherwise unperturbed. Once growth is attenuated by pharmacological inhibition of Ras signaling, this interaction is gone.

In contrast, a negative differential interaction occurred between *Rel* and *pnt*. While *Rel* knockdown rescued the fitness-defect induced by *pnt* knockdown under normal conditions, it aggravated the *pnt* knockdown phenotype after MEK inhibition (Figure 4D). Thus, we hypothesize that both, the aggravating interaction between *skd* and *Stat92E* and the alleviating interaction of *Rel* and *pnt* depend on the proper function of Dsor1. Mixed forms, such as interactions that deviate strongly in the beginning of an experiment and converge later, were also observed.

To assess whether different features (grouped to meta features) or pathways show enrichments in one or the other interaction type, we analyzed enrichment of interaction counts over a random distribution. We found considerably more stable than differential interactions for all feature (18468 vs. 4723, 16 phenotypic features, Figure 4E). While the distribution of negative and positive interactions over all features was symmetric, selected phenotypic features capture high numbers of alleviating (nuclear shape) or aggravating (nuclear texture) stable interactions.

We next sought to validate the linear models to describe time and treatment dependent genetic interactions. To this end, we compared the residuals of each fit with the actual experimental variance measured at each time point. If the model fails to fit the data appropriately (e.g. the comparison does not behave monotonic) one would expect that the residuals are unexpectedly greater than the variance. However, analyses of all interactions for each phenotypic feature reveals that this is rarely the case (Figure 4F, F’). In most models, remaining residuals of fit are explained by the variance between biological replicates (avg. PCC = 0.96, R^2^=0.92). We thus concluded that the π-score dependence on time and treatment can be reliably quantified using linear model statistics.

Additionally, we found that differential interactions, compared to stable interactions, enriched in specific signaling pathways related to MEK inhibition. While, for example, ribosome or spliceosome related genes formed mostly alleviating and condition stable interactions (Figure 4G), the JNK pathway was enriched for alleviating stable and negative differential interactions (Figure 4H). Other pathways, such as Ras signaling, *Rel*, Mediator signaling or Jak/STAT signaling were equally overrepresented in stable and differential interactions. Among the pathways tested, the enrichment of stable interactions highlights pathways which large impact on the interaction network controlling cell viability. The enrichment of differential interactions highlights pathways that are sensitive to MEK inhibition.

Differential genetic interactions are not equally distributed over all genes that were tested. Jak/STAT signaling components (*Stat92E, dome, upd3*) alongside Ras signaling members (*drk, rl, dos, Sos, pnt*) and, interestingly, Imd signaling (Rel) showed specific enrichment of differential interactions (cell count feature, Figure 4I). Specifically, *pnt* forms many positive differential interactions (alleviated upon MEK inhibition) while *Pvr* is involved in many negative differential interactions (aggravated by MEK inhibition). This could be attributed to *pnt* acting as a terminal transcriptional effector of the signal triggered by the activated receptor *Pvr*. We also found that genes, which form more stable genetic interactions also enrich differential interactions (compare linear trendline, Figure 4J). However, some particular genes are involved in unexpectedly many differential interactions. This indicates that a rather specific response to the treatment is reflected in the differential interactions. These data demonstrate that time dependent modeling of interaction scores sensitively detects treatment differential interactions which enrich in and thus highlight Ras sensitive biological processes.

### Signaling pathways rewire with different time dependencies

MODIFI estimates the time dependence (σ) of each differential interaction. This term can be interpreted as the slope by which an interaction changes (e.g. strengthens or weakens) over time. Depending on the initial difference (compare Figure 3A), π-scores increase or decrease over time, diverge or converge. The most abundant interaction in this study describes a treatment invariant interaction that could not be measured initially but forms over the course of the experiment (78 % of all significant interactions, FDR<0.1).

In the following analyses, we use genetic interactions based on cell count as an example to test whether genes or pathways react at different specific rates. For example, from 48 h to 96 h after compound addition, genetic interactions with *Rel* remained stable, whereas interactions of Jak/STAT or Ras signaling-related genes changed significantly over time. Interactions with housekeeping related genes (proteasomal or ribosomal subunits) show phenotypes of an exceptionally high time dependence (Figure 5A). These data indicate that interactions of the different biological processes rewire at different rates after perturbation.

**Figure 5.**
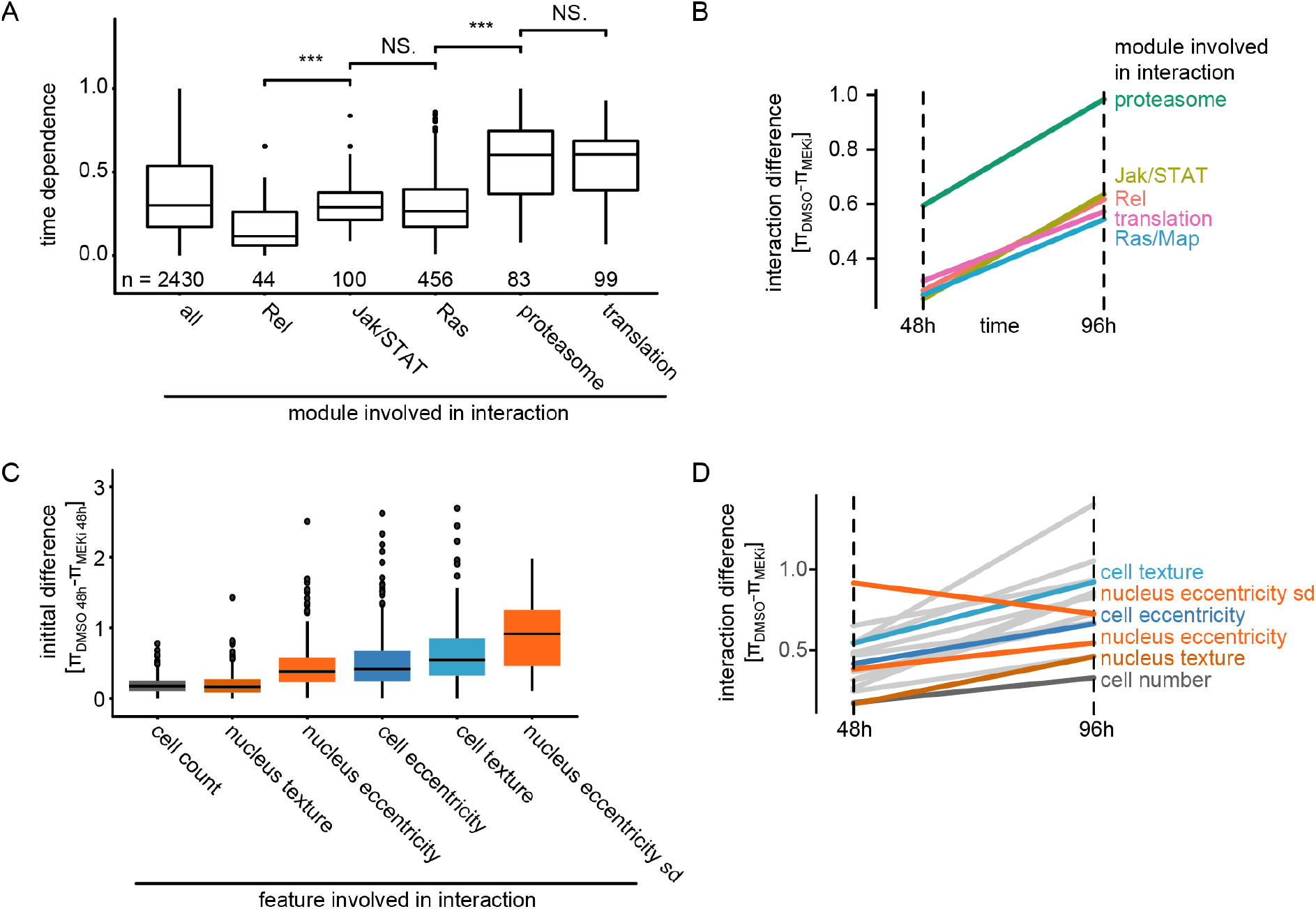
Timing and initial difference of interactions depend on the biological process and feature. (A) Time dependence (σ) of interactions stratified by biological process. Boxplots show the median (black bar), the 25^th^ and 75^th^ percentile (box) ± 1.5 times the interquartile range (whiskers). Points outside that range are plotted individually. Significance is tested by a two-sided welch t-test (***: p<<0.001, NS: p>0.05). Data is shown based on significant (FDR < 0.1) cell count based interactions involving genes belonging to this process. (B, B’) Median π-score differences stratified by pathway annotation of affected genes. All significant (FDR<0.1) time dependent interactions based on cell count feature are summarized by median. Interactions formed by genes that are proteasome associated show the highest initial difference and steepest increase over time. (C) Initial difference of interaction scores 48 h after treatment stratified by feature. Boxplots show the median (black bar), the 25^th^ and 75^th^ percentile (box) ±1.5 times the interquartile range (whiskers). Points outside that range are plotted individually. All features shown (except nucleus texture) show significantly (p>>0.001, two-sided student’s t-test) higher initial differences than cell count based interactions. (D) Median π-score differences for the first and the last measured time point. Trajectories for all features are shown over all genes that showed a significantly time dependent interaction (FDR<0.1). Features are highlighted by their feature group. All features except nucleus eccentricity measure interaction differences that become more profound over time.

We also hypothesized that the difference of interaction scores and its time dependence could inform about the influence of MEK inhibition on different biological modules or phenotypic features. Cell fitness-based interactions formed by proteasome related genes show the strongest phenotypic differences between treatments at the initial and last measured time point (Figure 5B). This suggests that proteasome related genes are involved in particularly strong differential interactions upon MEK inhibition. These interactions interfere with cell proliferation early on during our experiment and also become stronger over time. This supports reports of synergistic effects between proteasome and MEK inhibition on perturbing cell viability (Leow et al., 2013).

Next, we hypothesized that phenotypic features measure different initial interaction differences and analyzed initial π-score differences between phenotypic features. Especially, cell morphology features (nucleus/cell eccentricity) and their variance within the population of cells show initial differences that are significantly (p<0.0001) higher than those measured by cell count (Figure 5C). Of note, nuclear eccentricity and its variance among the population of cells (nucleus eccentricity sd) are also the only initially different features that are masked later on. All other phenotypes show an increased interaction difference over time (Figure 5D). Cell count shows the smallest interaction differences between the treatments in general, irrespective of the time point. Together, these analyses demonstrate that the time dependence of genetic interactions is specific to certain biological process. It further highlights that phenotypes beyond cell viability capture early differential interactions.

### A correlation network of differential interactions maps genes into functional modules

Next, we analyzed whether interaction networks formed by different biological modules or core signaling pathway change systematically over time and treatment. In the following examples we used cell eccentricity as an exemplary feature which we found to capture early cellular responses. Figure 6A shows how an interaction sub-network including Jak/STAT signaling, Ras/Map signaling components and spliceosome related genes rewires over time in reaction to MEK inhibition. Core housekeeping modules (ribosome, spliceosome or proteasome) were highly interconnected by alleviating stable interactions. In contrast, components of the Ras signaling, Jak/STAT signaling or Tor signaling cascade showed aggravating interactions with housekeeping modules. We observed that (i) alleviating interactions (π>0) dominate early time points, (ii) many initially alleviating interactions reverse over time (π>0 → π<0), (iii) differences attributed to the compound treatment become more profound over time. Lastly, we noted that genes in proximity tend to have similar interaction patterns coherently changing over time and treatment (**Supplementary File 1**). Previous studies implied that similarities of differential genetic interaction profiles can identify functionally related genes (Bean and Ideker, 2012). Thus, interactions of related genes change coherently upon network perturbation. Hence, we defined differential interaction profiles for each target gene. We used the modeled interaction difference between treatments over time (δ) to quantify interaction change due to Dsor1 inhibition. For every target we calculated δ with every query gene in a vector comprising 76 measurements for cell eccentricity.

**Figure 6.**
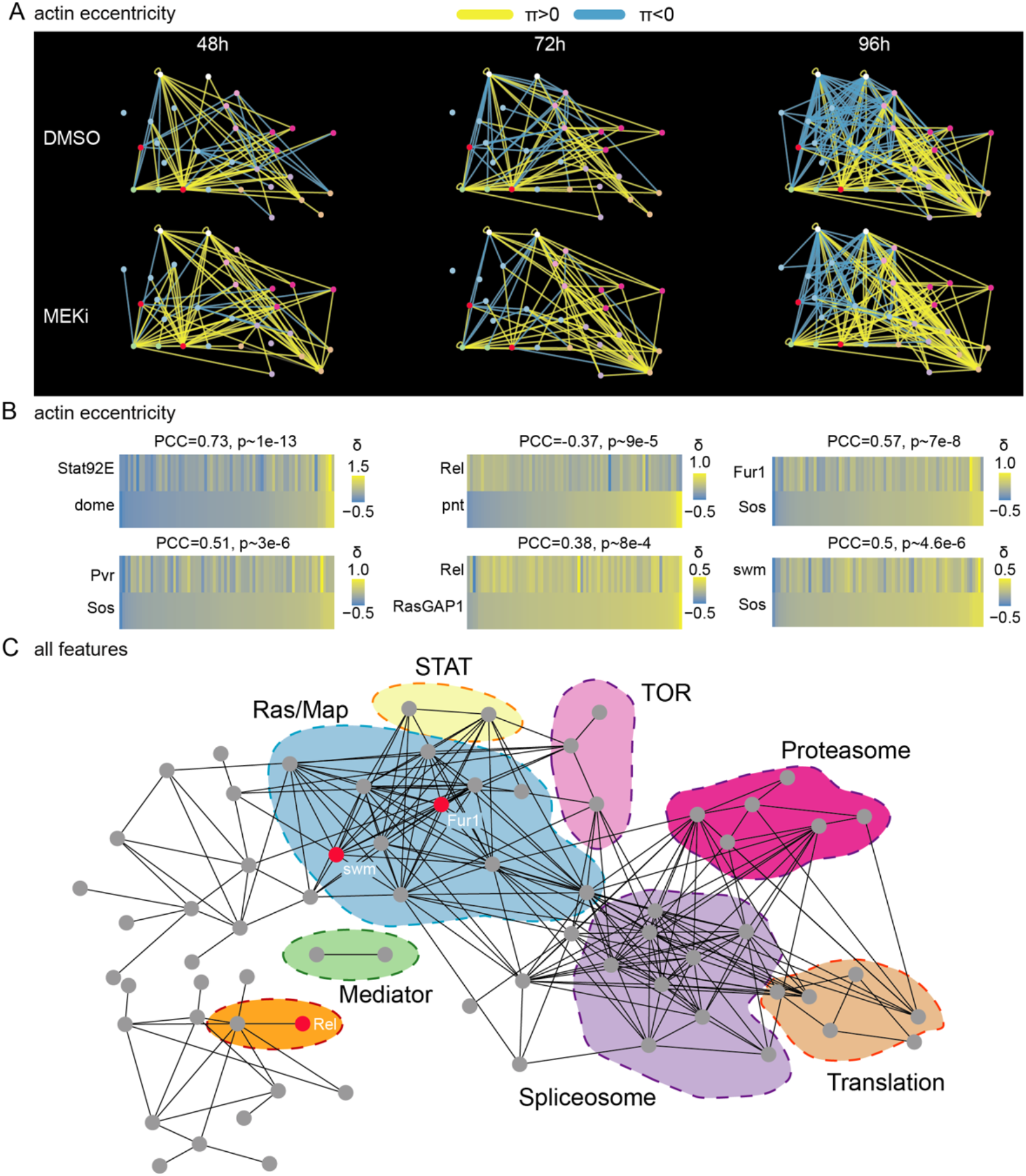
A correlation network of differential interactions maps pathways modules. (A) Network of genetic interactions between selected genes. The networks include all candidate Ras-signaling (blue), Jak/STAT-signaling (white), Tor-signaling (pink), proteasome (red), translation (orange) and splicing (purple) related genes. Significant (FDR<0.1) alleviating interactions are shown in yellow. Significant (FDR<0.1) aggravating interactions are shown in blue. All interactions are based on the cell eccentricity feature. Interactions become more abundant and stronger over time, more alleviating interactions can be observed under MEK inhibition. Ras and Jak/STAT related genes are mostly connected by aggravating interactions. (B) Correlations of differential interaction profiles between known regulators and candidate genes. Profiles of all δ-scores along cell eccentricity and 76 query genes were constructed for 168 target genes and pairwise correlations were calculated. Shown is the Pearson correlation coefficient (PCC) and asymptotic p-value as implemented in the R package Hmisc. All correlations shown are significant with an p-value<0.01. Jak/STAT and Ras components show high correlations as expected. *Rel* appears as negative, *Fur1* and *swm* as positive regulators of Ras signaling. (C) Pairwise correlation network of differential interaction profiles across all 16 features. Shown are all genes with at least one edge. Edges are drawn if two gene’s δ-profiles correlate with PCC>0.5. Nodes are ordered by force directed spring embedded layout. A high degree of clustering of known pathways indicates meaningful correlations.

Correlations between profiles (Figure 6B) confirmed known functional relationships of genes, as for example the profiles of the genes *Stat92E* and *dome*, members of the *Drosophila* Jak/STAT pathway, were similar (PCC 0.73) confirming that both genes share biological function upon perturbation of Ras signaling (Xu et al., 2011). Furthermore, our analysis showed a correlation of differential genetic interactions for all features between *Stat92E, dome* and Ras signaling. Interestingly, the profile of *Rel* was similar to negative regulators of Ras signaling *(RasGAP1*, PCC 0.38), but was anticorrelated with positive regulators *(pnt*, PCC −0.37) indicating a potential crosstalk between the two pathways.

We expected that a correlation-based network drawn from differential interaction profiles across all phenotypic features reveals modules of functionally related genes. Thus, we calculated the pairwise correlation coefficients (PCC) of differential interaction profiles (interactions with 76 query genes) including all 16 cellular features of all 176 target genes. We visualized resulting positive correlations in a network graph highlighting biological processes and candidate genes (Figure 6C, **Supplementary File 2**). This revealed that correlations of differential interaction profiles clustered genes into known pathway modules. Of note, *Rel* and *Fur1* (*FURIN*) and *swm* (*RBM26*) showed unexpected correlations with members of the Ras signaling cascade (Figure 6B).

It is expected that genes with similar tasks irrespective of the treatment show similar interaction profiles between and within conditions. In contrast, genes with a treatment dependent function should lose or gain correlations to other genes when compared between treatments (Billmann et al., 2017). To test this, we defined profiles of all interactions across all cellular features and time points and correlated them between genes and between conditions. Most interaction profile correlations did not differ significantly between conditions, compared to within conditions (**Supplementary Figure S7**). Specifically affected gene pairs were mostly Ras signaling components. Interestingly, also profile correlations of Jak/STAT signaling components (*Stat92E, dome*) as well as of the two genes *Fur1* and *swm* differed between and within conditions. This provides further clues that *Fur1* and *swm* are implicated Ras signaling. Only few, weak interaction profile correlations were higher between than within conditions.

### Rel and pnt act in a MEK-dependent negative feedback loop

We have shown that the differential interaction profiles of *Rel* and pnt were negatively correlated, whereas *Rel* profiles were positively correlated with *RasGAP1*, a negative regulator of Ras (Figure 6B). This suggested that *Rel* itself might function as a negative regulator of Ras signaling. We observed that *Rel* depletion alone had little impact on cell growth, as compared to *pnt*, but showed a cell length (major axis) phenotype (Figure 7A, **Supplementary Figure S4**). Co-depletion of *pnt* and *Rel* altered both cell number and cell length. Under control conditions depletion of *Rel* alleviated the loss of viability and cell length phenotypes after *pnt* knockdown (Figure 7A). This interaction was attenuated under MEK inhibition (Figure 7B) when co-depletion of *Rel* and *pnt* led to a synthetic lethal phenotype (FDR<0.1, Figure 7C, C’). These interactions were observed for both dsRNA designs (PCC=0.88 and 0.96 for *Rel* and *pnt*, **Supplementary Figure 8**).

**Figure 7.**
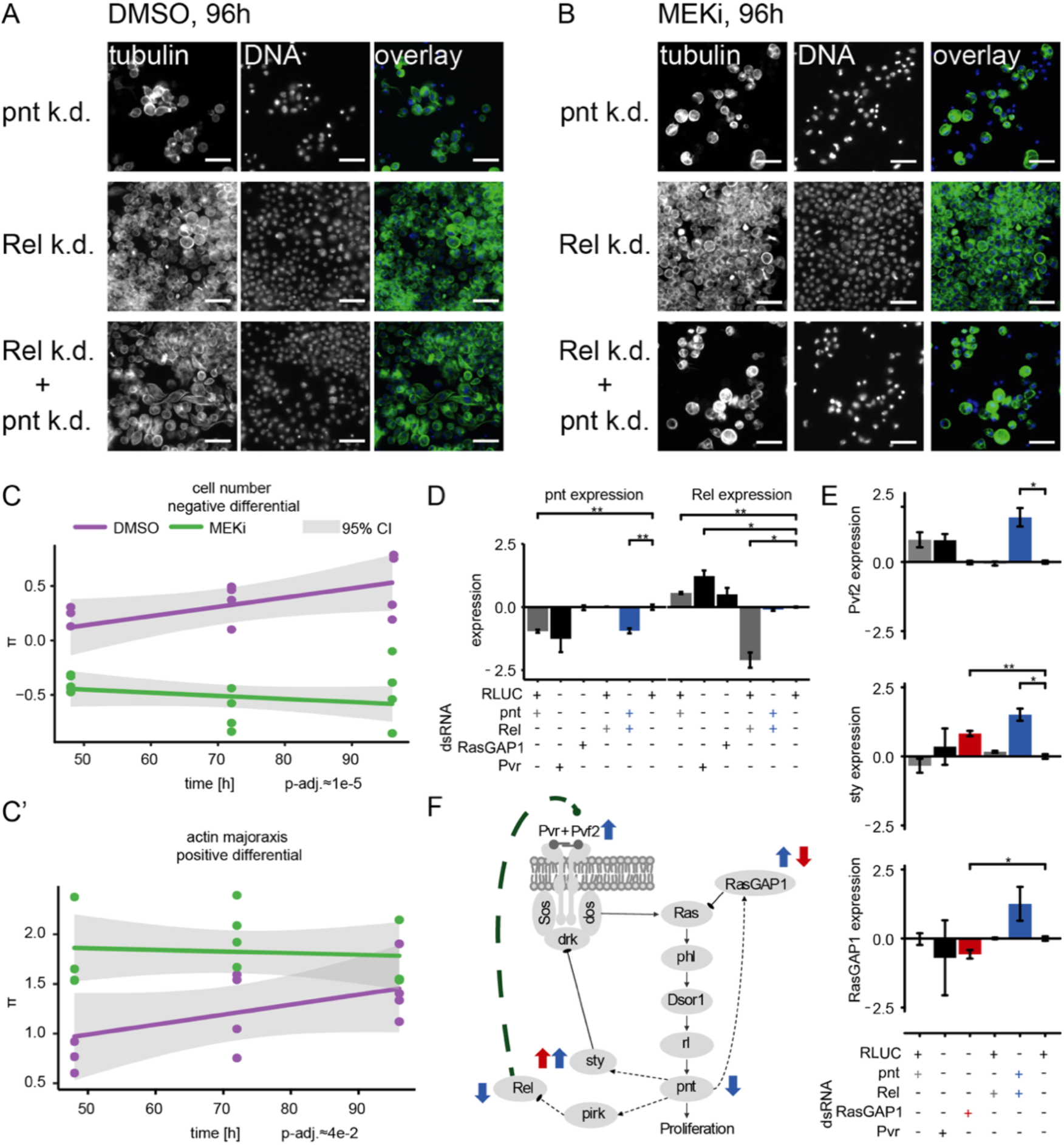
Crosstalk of NF-κB and Ras signaling through *Rel* and *pnt*. (A) Upon *Rel* or *Rel/pnt* knockdown cells behave normally, growth is inhibited upon *pnt* knockdown alone. (B) Dsor1 inhibitor treatment attenuated this alleviating interaction. Scale bar=30 μm. Images are pseudo colored, DNA/DAPI = blue, FITC/α-tubulin = green. (C) Quantified negative differential interaction between *Rel* and *pnt*. The trajectory of the Dsor1 inhibitor treatment is lower than the solvent control condition for cell count interaction indicating synthetic lethality under MEK inhibition. (C’) Actin major axis shows a strong positive interaction (cells are enlarged like under *pnt* knockdown). Error of fit is shown as 95 % confidence interval. Significance of model fit was analyzed using the Wald test on the robust linear model (R/sfsmisc/f.robftest). (D, E) Expression of candidate and marker genes assessed by qPCR (3 days RNAi treatment, n=3, log_2_ fold_RLUC_, mean ± s.e.m., t-test) on Dmel2 cells. (D) *pnt* expression is reduced upon *pnt* and *Pvr* knockdown. *Rel* knockdown does not rescue *pnt* expression. *Rel* expression is increased upon *pnt* and *Pvr* knockdown and decreased upon *Rel* knockdown. Upon *pnt* and *Rel* knock down, *Rel* expression is rescued to normal levels. (E) Pvf2 expression is induced only upon *Rel/pnt* double knockdown. This leads to increased expression of sty and *RasGAP1. RasGAP1* knockdown increases sty expression and decreases *RasGAP1* expression. (D-E) *~p<0.05, **~p<0.01. (F) A model summarizes the qPCR results in context of the Ras signaling cascade. Dashed lines are transcriptional interactions, solid lines are protein-protein interactions. All black interactions are known, while the green interaction is inferred from the data. Blue arrows indicate that *Pvf2, sty and RasGAP1* were up-regulated upon *Rel/pnt* co-knockdown and by that Ras pathway activity was restored. A similar pattern could be observed upon *RasGAP1* knockdown, which causes intrinsic hyperactivation of Ras signaling by constitutive Ras activation (measured by upregulation of *sty*, red arrows).

*Pvf2* (orthologue of human VEGF) is up-regulated in the absence of *Rel* (log_2_ fold-change~1.5) (Boutros et al., 2002). The data presented here indicate that a knockdown of *Rel* induced a re-activation of the Ras pathway which is dependent on Dsor1 activity (Figure 7A-C). We hypothesized that *Rel* negatively regulates Ras signaling by repressing the expression of *Pvf2*, the ligand activating the Pvr-Ras-phl-Dsor1-rl-pnt signaling cascade after binding to Pvr (PDGFR). To test this hypothesis, we performed qPCR analysis of *pnt*, *Rel*, *Pvf2*, *sty* (*SPRY2*) and *RasGAP1* expression levels (Figure 7D, E, **Supplementary Figure 8**). We first confirmed the up-regulation of *Rel* after depletion of Ras (Figure 7D) and showed that up-regulation of *Rel* was suppressed by *pnt* co-RNAi. *Pvr* knockdown, as a control for loss-of Ras signaling activity, led to a downregulation of *pnt* and *RasGAP1. Pvr* knockdown also induced a strong upregulation of *Rel* expression. Finally, co-RNAi of *Rel* and *pnt* induced a significant increase in *Pvf2* expression, not observed by depletion of either gene alone (Figure 7E). The *Rel/pnt* co-RNAi also induced upregulation of negative regulators of Ras signaling *sprouty* (*sty*) (Casci et al., 1999) and *RasGAP1* (Feldmann et al., 1999) (Figure 7E), thereby providing a mechanistic explanation how *Rel* could negatively regulate Ras signaling.

We hypothesized that this regulatory loop is mediated by the transcriptional regulation of *Pvf2* and requires Dsor1 mediated Ras signaling activity, as summarized in Figure 7F. These changes were observed both at 48h and 96h time-points (**Supplementary Figure 9 and 10**). Interestingly, protein levels of rl were down regulated by *pnt*-or *rl*-RNAi and rescued by *Rel* co-RNAi (**Supplementary Figure 10F, G**). Overall, these experiments provide a mechanistic basis how *Rel* acts as a negative regulator of Ras signaling in a context-dependent manner.

## DISCUSSION

To better understand context-dependent differences in genetic networks upon changes in environmental conditions is a current frontier in genetics (Rancati et al., 2018). Many biological processes rely on context-dependent changes in genetic requirements, from robustness of cell differentiation during development to responses of cancer cells to chemotherapeutic treatments. However, only few studies on selected phenotypes have systematically analyzed how environmental changes impact genetic interaction networks. Previous studies have analyzed genetic networks after activation of the DNA damage response signaling in yeast or changes in Wnt signaling activity in *Drosophila* cells (Bandyopadhyay et al., 2010; Billmann et al., 2017; Díaz-Mejía et al., 2018). In these studies, positive and negative differential, and stable interactions have been determined based on fitness phenotypes or pathway reporter activity in static end-point assays. Aim of the present study was to analyze changes in genetic networks that impact a broad spectrum of phenotypes by imaging and multiparametric image analysis and to determine how differential interactions change over time after small molecule perturbation of the Ras signaling pathway.

To this end, we established a high throughput image-based assay which enabled us to reproducibly measure many phenotypes including cell proliferation and cell morphology which are influenced by many cellular processes (Breinig et al., 2015; Fuchs et al., 2010; Horn et al., 2011). We used this assay to measure genetic interactions between differential treatment conditions over the course of three time points. To this end, we assessed the phenotypes of 76608 di-genic interactions in *Drosophila* hemocyte-like cells in total. Each interaction was characterized by a vector of 16 non-redundant and reproducible phenotypic features. Further, we developed MODIFI, a two-factor robust linear model to quantitatively describe the time and treatment dependent changes of genetic interactions. MODIFI also allowed us to describe whether an interaction is differential between treatments (treatment could predict π-score) or time dependent (time predicted the π-score).

Our analysis showed that we detected more differential interactions as compared to endpoint measurements or single time point replicates. Enrichment of differential interactions among stress responsive pathways and genes underlines their biological relevance. Using this approach, we also analyzed the treatment (δ) and time (σ) dependency of interactions of specific genes and pathways. Overall, we found that differential interactions were called with much higher confidence when comparing treatment dependent phenotypes along time. Furthermore, the measurement of multiple phenotypic features at the same time enabled more detailed characterization of the observed differential interaction. We also tested whether the establishment of phenotypes is dependent on a gene’s expression level but found no correlation of high gene expression and high time dependency (data not shown). Our data further suggests that σ is influenced by the general resilience of a pathway or signaling module to perturbations. This makes it unlikely that the stability or turnover of a single gene’s product is a major driver of time-dependent establishment of genetic interactions. We found, for example, that genetic interactions of ‘core’ (or housekeeping) modules such as the translation machinery, proteasome and others induce phenotypes that are much stronger at later time-points in the experiment. In contrast, intra-cellular signaling modules such as signaling and innate immunity ‘rewire’ early in the experiment. Our analysis classified genes into categories of genetic interactions that are (i) signaling modules central to the cells’ physiological role, (ii) signaling modules required for maintaining homeostasis and (iii) resilient ‘core’ modules whose network hubs form interactions on a longer timescale. We also found that measuring different phenotypes provided more information about the development of interaction differences over varying time scales. While the cell count (comparable to yeast colony size) as a phenotype captures cellular reactions rather late in the experiment, other phenotypes, such as nuclear morphology or cytoskeleton texture, enabled to measure immediate cellular reactions.

In the chemico-genetic experiments, we found that pathways interacting with Ras signaling reacted strongest to the Dsor1/MEK inhibition. To map signaling modules that react similarly towards Ras signal perturbation, we correlated δ-profiles along all features between all target genes. By this means, genes whose interactions change coherently upon Dsor1 inhibition are grouped into highly inter-connected modules. Consequently, this correlation network clusters genes of similar functions in proximity with each other. Each module is also characterized by a coherent reaction towards Dsor1 perturbation. Interestingly, *rolled (rl), Dsor1* and *pole hole* (*phl*) (ERK1/2, MEK1/2 and Raf) were not connected to the rest of Ras signaling related genes in the correlation network. In contrast, they correlated with Ras when only using interaction profiles of the control treatment, indicating that the chemico-genetic analysis identified ‘responsive’ factors that can be uncoupled upon environmental modulation of specific signaling modules.

Our analysis also revealed three genes that unexpectedly connected to Ras signaling: *Fur1*, a serine-type endopeptidase (Kim et al., 2015), swm, involved in mitotic checkpoint regulation and hedgehog signaling (Casso et al., 2008; Dong et al., 1997) and *Rel* (Foley and O’Farrell, 2004). The correlation of *Fur1* and *swm* with positive regulators of Ras signaling indicates that they respond similarly towards Dsor1 inhibition as Ras pathway members. In addition, we identified *Rel (NF-κB)* as a strong differential genetic interactor, suggesting that mitogenic Ras signaling and innate immune pathways are interdependent. Once *Rel* is lost, cells become more dependent on Ras signaling; a phenotype that can be blocked by perturbing Dsor1 activity chemically or genetically. Already at a low dose, both interferences result in a synthetic lethal phenotype that kills *Drosophila* hemocyte-like cells. Conversely, it was previously shown that Ras signaling influences *Rel* activity by regulation of its negative transcriptional regulator pirk (Ragab et al., 2011). We hypothesize that this mutual negative feedback regulation could be the basis for a ‘fight’ or ‘flight’ response of the immune cells; balancing an immune and proliferative response in the same cell.

Large-scale studies on gene essentiality have challenged the concept of a static repertoire of essential genes. In contrast, loss-of-function screens in different genetic background of cancer cells identified ‘core’ and ‘genotype’-dependent sets of essential genes. This indicates that essentiality is modulated in a context-dependent manner (Hart et al., 2015; McDonald et al., 2017; Rauscher et al., 2018). At this point, our study is the largest exploration of gene-gene-drug interactions based on multiparametric, non-essentiality phenotypes. We demonstrate how different vulnerabilities for a diverse set of automatically scored phenotypes change upon time and environmental conditions. Our modeling approach increases the confidence to call differential interactions upon changes of environmental conditions. This allows to map a correlation network of cellular modules that react coherently towards the external stimulus. We expect that, when further studies of context-dependent genetic interactions will be become available, a comparative analysis will provide fundamental insights into how different cellular networks react to environmental stimuli with implications for therapy resistance and timing of drug treatments. This study introduced an experimental and analysis framework to explore time-dependent rewiring of genetic networks which can be used to dissect the complexity of biological networks in model organisms and human cells.

## METHODS

### Genome-wide RNAi library

We used a genome-wide *Drosophila melanogaster* dsRNA library (HD3) in this study, as previously described (Billmann et al., 2017; Horn et al., 2011). The library contains 28941 dsRNA reagents targeting 14242 unique gene IDs in the *Drosophila melanogaster* genome and contains two sequence independent reagents targeting 13617 IDs twice and the remaining genes once. The reagents were optimized for the BDGP5 mRNA annotations in *Drosophila melanogaster* by for example avoiding CAN repeats and non-unique sequences (off-targets). 250 ng dsRNA, synthesized as described previously, were aliquoted to 384 Greiner μClear plates prior to the image-based assay at a mass of 250 ng/well. A table containing all sequences that were used in the genome-wide RNAi screen can be found in **Supplementary Table S5**. Another table containing sequence IDs (HD3) that were used in the combinatorial RNAi screen can be found in **Supplementary Table S6**.

### Image-based RNAi screening

dsRNA reagents dissolved in water were spotted into barcoded 384-well microscopy plates (Greiner μ μClear, black, flat-transparent-bottom, Ref: 781092, Greiner Bio One International GmbH, Frickenhausen, Germany) to reach a final mass of 250 ng dsRNA per well (5 μl of a 50 ng/μl solution). Express V medium (Gibco, Ref: 10486-025, Life Technologies GmbH, Darmstadt, Germany) with 10 % Glutamax (Gibco, Ref: 35050-061) was pre-warmed to 25 °C and 30 μl were dispensed on top of the spotted dsRNA using a MultiDrop Combi dispenser, standard cassette (Thermo Fisher Scientific, Ref: 5840400, Life Technologies GmbH, Darmstadt, Germany).

10 μl of pre-diluted Dmel2 cell suspension were seeded to a final concentration of 9000 cells/well into the prepared assay plates using MultiDrop Combi dispensing under constant stirring of the suspension in a sterile spinner flask (Corning, Ref: CLS4500500, Kaiserslautern, Germany). After cell addition, the assay plates were heat sealed using a PlateLoc (peelable seal, 2.3 sec at 180 °C, Agilent Technologies Deutschland GmbH & Co. KG, Waldbronn, Germany) and centrifuged at 140x g for 60 sec. Cells were incubated for 24 h at 25 °C without CO_2_ adjustment.

After 24 h incubation, plates with growing cells were opened and small molecule treatment was performed. The concentration of applied compound is outlined with the separate experiments in the following paragraphs. Per well 5 μl of a solution containing 5 % DMSO (Sigma Aldrich, Ref: 41644-1l, Merck KGaA, Darmstadt, Germany) in medium, or PD-0325901(Cayman chemical, Ref: CAY-13034-5, Biomol GmbH, Germany) dissolved in 5 % DMSO in medium, were added to achieve a final assay concentration of 0.5 % DMSO and varying small molecule concentrations. We found that the most drastic phenotypic changes among a number of features (examples shown in **Supplemental Figure 1G**) occurred in a concentration window around the drug’s ED50 (1.5 nM, GFP dsRNA treatment was paired with compound treatment). Thus, we selected a concentration of 1.5 nM PD-0325901 as an optimal condition for the following screening experiments, ranging within an order of magnitude of the ED50 known for treatment of mammalian tissue cells cultures (Ciuffreda et al., 2009; Hatzivassiliou et al., 2013). After compound addition, plates were sealed again and incubated at 25 °C without CO_2_ adjustment for 48 h, 72 h or 96 h depending on the experiment.

Assays were stopped after the second incubation period by fixation using a robotics procedure on a CyBioWell vario (384-well pipetting head, Analytic Jena AG, Jena, Germany). There, supernatant was removed, and cells were washed with 50 μl PBS (Sigma Aldrich, Ref: P3813-10PAK) per well. After addition of 40 μl Fix-Perm solution (4% Para-formaldehyde (Roth, Ref: 0335.3, Karlsruhe, Germany); 0,3% Triton X-100 (Sigma Aldrich, Ref: T8787- 250ml); 0,1% Tween20 (Sigma Aldrich, Ref: P1379-100ML); 1% BSA (GERBU Biotechnik GmbH, Ref: 1507.0100, Heidelberg, Germany), plates were incubated for 60 min at RT and then washed twice with 50 μl of PBS. 50 μl of PBS were added again and plates were stored at 4 °C before staining. For staining, remaining PBS was removed and fixed cells were first blocked by adding 30 μl of blocking solution (4% BSA; 0,1% Triton X-100, 0,1% Tween20) and incubated for 30 minutes at RT. Next, the blocking buffer was removed and 10 μl of staining solution (1:4000 Hoechst (Thermo Scientific, Ref: H1399, Life Technologies GmbH, Darmstadt, Germany), 1:1500 primary FITC labelled anti α-tubulin antibody (Sigma Aldrich, P1951), 1:6000 Phalloidin-TRITC conjugate (Sigma Aldrich, F2168-.5ml) in 1x blocking buffer) were added. After addition of the staining solution plates were incubated for 60 min at RT. After staining, 30 μl of PBS were added and the staining solution was removed. After two additional washing steps with 50 μl PBS another 50 μl of fresh PBS were added per well and stored at 4 °C until imaging.

### Genome-wide chemico-genetic interaction screening

We performed genome-wide RNAi screens in combination with drug and solvent control treatment to verify dsRNA reagent efficiency, identify candidate genes for combinatorial screening and to find which genes react most differentially to the *Dsor1* inhibitor (PD-0325901) treatment. Four sets of 88 x 384-well Greiner μClear plates were spotted with the HD3 library, 5 μl of 50 ng/μl dsRNA in each well. The HD3 library is arrayed to target one gene with one dsRNA design per well. Two additional plates, containing only controls were added to control assay reproducibility, robustness and effect size. Controls were chosen to spread over the complete dynamic range of cell fitness. dsRNAs against *RLUC* and *GFP* expressing plasmids serve as non-targeting negative controls, such that we could control for unspecific dsRNA induced phenotypes. dsRNA containing plates were thawed, seeded with cells and left for 24 h at 25 °C without CO_2_ adjustment for incubation prior to drug treatment. Plates were opened and 5 μl of 15 nM PD-0325901 in 5 % DMSO were added resulting in a final assay concentration of 1.5 nM PD-0325901 in 0.5 % DMSO in medium. Cells were left to incubate for another 72 h at 25 °C without CO_2_ adjustment prior to fixation, staining and imaging. Images were acquired using the standard protocol described below with low illumination timings (DAPI: 100 ms, Cy3: 200 ms, FITC: 300 ms). The resulting images were analyzed in line with the acquisition using the standard image analysis pipeline and progress was monitored using our automated analysis pipeline as described below.

### Combinatorial RNAi screening under differential time and treatment conditions

The design of the library for combinatorial screening is described in a separate paragraph. 168 genes were chosen for design of a combinatorial RNAi library. The dsRNA sequences that were used in the combinatorial library can be found in **Supplementary Tables S5 and S6**. All used labware and reagents, which are not further detailed here have been the same as in previous experiments. The library contained 12 batches for screening, each comprising 80 x 384-well Greiner μClear plates spotted with 250 ng dsRNA/well dissolved in 5 μl of DNase, RNase free water. dsRNAs were obtained from the HD3-library templates and synthesized accordingly. To avoid contaminations, all dsRNAs were sterile filtered using Steriflips-0.22μm (Merck Millipore, Ref: SCGP00525, Darmstadt, Germany) for the query dsRNAs and MultiScreenHTS-GV 0.22 μm filter plates (Merck Millipore, Ref: MSGVN2250) for the target dsRNAs. Genes were divided into target and query genes based on prior knowledge on key pathway components and screened a matrix of 76 genes combined with 168 genes. All query genes were included in the list of target genes. We screened each target gene in two sequence independent designs and each query gene in one design. This way, we screened 25536 dsRNA combinations (12768 gene pairs) in each batch. Combinatorial dsRNA spotting was achieved with combining the query and target master plates such that 2.5 μl of each query dsRNA were spotted onto 2.5 μl target dsRNA using a Beckman FX robotic liquid handling station (Beckman Coulter, Krefeld, Germany). In order to control for RNAi induced phenotypes and per-plate batch effects control dsRNAs against *Dsor1, drk, Diap1, RasGAP1, Pten, pnt, Pvr, Rho1* and *RLUC* expressing plasmid were spotted on each plate and not paired with a second query dsRNA perturbation. Two control plates containing only the target gene dsRNA reagents with 250 ng dsRNA per well complete one screening batch of 80 plates and controlled for screening batch effects. To achieve differential treatment and time resolution, 12 screening batches were prepared. They were divided into two groups of six batches, which then were treated under different conditions in duplicate. 6 library batches are needed to screen two conditions (1.5 nM PD-0325901 and 0.5 % DMSO) at three time points (fixation 48 h, 72 h, 96 h after small molecule addition), all in all comprising 480 screened plates. The entire experiment was repeated twice. This way we screened 960 x 384-well plates.

The assay workflow followed the same procedures as outlined above. Briefly, 9000 cells per well were seeded onto 384-well Greiner μClear plates for microscopy, which were prespotted with a combinatorial dsRNA library. After centrifugation, plates were sealed and left to incubate for 24 h at 25 °C prior to compound addition. Therefore, a PD-0325901 dilution (15 nM in medium with 5 % DMSO), and a 5 % DMSO-only dilution in medium were prepared and added to the opened plates. This resulted in either 1.5 nM PD-0325901 or 0.5 % DMSO in-assay concentrations. Plates were sealed again using a heat sealer and left to incubate until the experiment was stopped by fixation after 48 h, 72 h and 96 h, respectively. Stained plates were imaged using an InCell Analyzer 2200 automated fluorescence microscope according to the protocol described above with 20x magnification, 3 channels per field and 4 fields per well. Resulting images were analyzed using the R/EBImage based pipeline described below.

### High-throughput imaging

All plates were imaged using the same protocol. There, an InCell-Analyzer 2200 automated fluorescence microscope (GE Healthcare GmbH, Solingen, Germany) with a Nikon SAC 20x objective (NA = 0.45) was used. The microscope was adjusted to scan Greiner μClear plates by setting the bottom height to 2850 μm and the bottom thickness to 200 μm and the laser autofocus function was applied to identify the well bottoms with attached cells. This Z-position was used for image acquisition in three fluorescence channels: Hoechst (excitation: 390±18 nm, emission: 435±48 nm) at 400 ms exposure (100 ms in dose response experiments and genome wide screens), Cy3 (excitation: 475±28 nm, emission 511±2 nm 3) at 300 ms exposure (200 ms in dose response experiments and genome wide screens) and FITC (excitation: 542±27 nm, emission: 597±45 nm) at 300 ms exposure (300 ms in dose response experiments and genome wide screens). Four fields of view were imaged per well at 20x magnification each representing a 665.60 μm x 665.60 μm area covered (approx. 20 % of total well area) by 2048 x 2048 pixels. For plate handling, the microscope was equipped with a KinedX robotic arm (PAA Scara, Peak Analysis & Automation Ltd, Hampshire, UK) allowing a fully automated image acquisition.

### Automated image processing and high-throughput image analysis

Plates were imaged and analyzed in batches of 40 plates and a custom pipeline allowed parallel image processing and analysis by bundling images of fields and channels of several wells. During imaging an automated pipeline scheduled the processing of image files for each field of view through the following analysis workflow, here described representatively for one field. Raw images of three channels with a size of 8.4 MB (16-Bit grey scale, 2048 x 2048 pixels) per image were captured with the InCell Analyzer 2200 software and saved as TIFF files on a server cluster for image processing and analysis. The image processing-and analysis pipeline covered two main blocks, first a sequence of pre-processing steps which was followed by extraction of phenotypic features from single cells. First, the images were read in and each channel was assigned to the subcellular structure that was selectively stained with the above described assay (Hoechst: DNA, Phalloidin-Cy3: F-actin, α-tubulin-FITC). To identify cell and nuclei boundaries, a duplicate image of each channel is ln transformed, scaled between 0 to 1 and smoothened by a Gaussian filter using a sigma of one. This reduced optical noise, improved the dynamic range and smoothened the image gradients for further segmentation by thresholding. For segmentation, the normalized actin and tubulin images were binarized by global thresholding. Second, the cell nuclei were identified by applying a local adaptive average threshold to binarize the DNA channel nuclei image and assigning objects. The resulting binary image was then subjected to morphological operations of opening and hull filling such that filled objects with smoothly roundish outlines result. Offsets for segmentation were varied if the channels surpassed certain thresholds. If more than 30 nuclei were counted per field, the objects were subjected to further propagation of nuclei objects into the an á priori defined cell body mask. Starting from the nucleus objects as seed regions, the cell bodies are segmented by propagating the nuclei objects into foreground area (Carpenter et al., 2006). This strategy allowed to identify single cells and corresponding nuclei as objects. Using the segmented object outlines as masks, features on each object and channel were calculated on the original image using the R/Bioconductor package EBImage (Pau et al., 2010).

Specifically, numeric descriptors for 5 feature classes are defined in the computeFeatures function from EBImage (**Supplementary Table S1**): (i) shape features which quantify the shape of cells and nuclei, (ii) basic features that describe the summary statistics, such as 5 % quantiles, of pixel intensity within the borders of the object, (iii) moment features that describe the spatial orientation of the objects, (iv) Haralick features derived from a pixel intensity co-occurrence matrix as texture descriptors (Haralick et al., 1973) and (v) social features such as distance to the first 20 nearest neighboring cells. Social features are derived by a k-nearest-neighbor search based on the geometric center points of single cells. Single cell data were stored and aggregated to well averaged data by calculating the trimmed mean (q=0.01) of all cells belonging to all fields of one well and its standard deviation.

### Data processing and normalization

For the analysis of the genome-wide chemico-genetic screens the following analysis strategy was pursued: feature data was collected in a data frame containing per-well aggregated values as trimmed mean and standard deviation. This data frame was then reformatted to a 4-dimensional data cube featuring the dimensions: feature, plate, well, screen. Per feature, the feature’s minimum value was added to each value prior to log_n_(x+1) transformation to approach the features’ histogram to normal distribution. Following transformation, each plate in each screen was normalized separately for each feature by B-Score normalization (Ljosa et al., 2013; Mpindi et al., 2015). The B-Score normalization centers and scales the data to be the residuals of the median polish divided by the median absolute deviation (mad) across all values of the plate and thus be symmetrically centered around zero and scaled in units of the mad. Here, 38 representative features were chosen based on their biological significance (our ability to refer them back to cellular phenotypes) and their biological reproducibility between the two mock (DMSO) treated replicate screens and their information content, as measured by added variance (**Supplementary Table S1**).

For the combinatorial screens the obtained data frame containing one row for each well and columns containing the measured features and well, plate and batch identifier as annotation columns. Data acquired from single cells was aggregated by calculating the trimmed mean (q=0.01) for each feature extracted in the respective well together with its standard deviation. This way, outliers, produced by over or under segmentation of cells, were mostly excluded from further analysis. Data was normalized by dividing each feature in each plate by the median of the non-targeting control wells (if that was not zero). Further, the values of each feature were transformed on a logarithmic scale using the generalized logarithm with c being the 3 % quantile of the features value distribution over all values (Caicedo et al., 2017; Fischer et al., 2015). For each feature, data was subsequently scaled and centered around 0 by using the robust Z-transformation, where the feature median is subtracted from each value and the result was divided by the median absolute deviation (x’=(x-median(x))/mad(x)). After that, all features were in normalized units of median absolute deviation from the median of that feature and normalized per plate. The normalized feature vector provided the basis to all further analyses.

### Candidate selection for combinatorial RNAi screening

The metrics used for judging the quality of dsRNA reagents and to assess the gene’s suitability for the combinatorial screen are summarized in **Supplementary Table S2**. For this purpose, several metrics have been deployed. Summarized, the applied metrics were used to assesses for each individual gene in the genome-wide HD3 library (i) the quality of the RNAi reagent, (ii) the effect size of the induced phenotypes under solvent control treatment as well as the differential effect size of the differential phenotype between small molecule treatment and control conditions, (iii) the quality of the target gene as a candidate for gene-gene-drug combinatorial screening. Effect size was quantified using the Euclidean distance (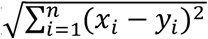) between sample and control measurements under different conditions. Quality of dsRNAs was assessed by calculating Pearson correlation coefficients between phenotypic profiles of biological and dsRNA design replicates. The quality of genes as screening candidates was assessed by gene expression analysis and literature analysis. The Q1 metric shows the strength of a knockdown induced profile when compared to the non-targeting control knockdown (here: GFP). This was calculated as the Z-Score normalized Euclidean distance of the genes profile to the control profile and can be used to inform if a phenotype of a gene is exceptionally strong or weak. In general, strong phenotypes (Q2) were preferred since they were more robust to experimental noise and are likely to engage in many genetic interactions (Costanzo et al., 2010). Q3 gives to what extend the phenotypic profile of those genes knockdown changes upon drug treatment. An ideal candidate for drug gene interaction screening shows a high value in this metric. Q4 and Q5 allow inferring the reproducibility of the measured phenotype by comparing the correlation of two sequence independent dsRNA designs targeting one gene and the correlation of one design across screen replicates, respectively. There, 7957 genes were targeted by designs whose feature vectors correlate with PCC > 0.5 while 17263 designs were reproducible between screens (PCC > 0.5). Q6 was used to infer if the respective gene is expressed under the screened conditions (Dmel-2 cells, 4 days in culture in Express-V medium). 12567 genes (88 % of all genes screened) had a log_2_ normalized read count greater than 0. In contrast, the knowledge sum in Q7 was used to avoid over enrichment of well characterized genes in the final combinatorial library. The “unknowne” was defined by means of assigning each gene a score describing how well it has been studied and characterized. Therefor the Gene Ontology terms associated to each *Drosophila* gene were downloaded from Flybase. In Flybase, each ontology term is annotated with evidence codes as provided by the gene ontology consortium (Ashburner et al., 2000). Each of these codes was then used to assign weights to the ontology terms for each gene (**Supplementary Table S7**). Ontology terms derived from experimental evidence, such as genetic interactions, direct assays or physical interactions were assigned the highest weight while computational annotations were weighted the lowest. For each gene, the sum of ontology terms was computed and used as a proxy for the current state of its functional characterization. For example, the cell fate determining receptor *Notch* is the most well studied gene with a score of 973, while all genes have an average score of 34.7 and the third quartile ends at 41. This means that only a minor fraction of genes is as well studied as *Notch* and most genes can be accounted as uncharacterized if their score is beneath 100 (90 % quantile). An example for such a gene is *tzn* with a knowledge sum of 14. Only known fact about *tzn* is its function as Hydroxyacylglutathione hydrolase in response to hypoxia (Alalouf et al., 2010; Jha et al., 2016). For screening, genes with a low knowledge sum were preferentially chosen.

### Modeling of genetic interactions

The data frame with normalized feature data per well was reformatted into a 5-dimensional data cube representing the experimental design. The dimensions are: target dsRNA (2 entries for each gene), query gene, time, treatment and feature. The data cube was further subjected to genetic interaction analysis following the protocol established by Bernd Fischer (Fischer et al., 2015; Horn et al., 2011; Laufer et al., 2013). There, genetic interactions are defined as the residuals of a modified median polish over the double perturbation matrix of one replicate, feature, treatment and time point. The median polish presents a robust linear fit (***M_ij_*** = ***m*_*i*_** + ***n_j_*** + **π_*ij*_** + ***ε***) that lifts the main effects (m, n) of each query such that it resembles the value of a single gene knockdown. The residuals of this fit scaled by their median absolute deviation are defined as π-scores. π-scores further provide us with a quantitative measure of genetic interaction following the multiplicative model plus some error term (ε) estimating the experimental noise. There, the interaction of two genes is defined as the deviation of the measured combined phenotype (M_ij_) from the expected phenotype for a target-query gene pair *i* and *j*. The expected phenotype is defined as the product of the two independent single knockdown phenotypes. The resulting π-scores are then collected for all replicates (dsRNA and experimental, each interaction is measured four times). The significance of their mean over all measured scores is estimated by a moderated students t-test as is implemented in the R-package limma. There, the t-test is adapted for situations where a small amount of observations is tested in many tests, normally causing large test variability, using an empirical Bayes variance estimator. P-values were adjusted using the methods of Benjamini Hochberg (Benjamini and Hochberg, 1995). From there on, adjusted p-values can be treated as false discovery rates. The FDR estimates the chance that the finding was observed by random chance given the entire dataset. This described procedure was applied to quantitatively calculate genetic interactions for each phenotypic feature.

### qPCR analysis

Quantitative real time PCR (qPCR) was used to analyze the transcriptional response following Rel/pnt co-RNAi. To this end, as 5*10^5^ cells / well were seeded in 630 μl ExpressFive (Gibco) culture medium and reverse transfected with 14 μg dsRNA. All dsRNAs denoted with #2 were used in 3 biological replicates and combinatorial RNAi was achieved by mixing 7 μg of dsRNA targeting each gene (**Supplementary Table S8**). After 72 h incubation (25 °C, no CO_2_ adjustment), cells were washed once in 750 μl PBS (Gibco) and lysed in 350 μl RLT buffer shipped with the RNAeasy-mini Kit (Qiagen). RNA was then purified from all samples according to manufactures standard instructions for spin column purification. An optional DNase digestion step was performed using the RNase-Free DNase Set (Qiagen). Samples were prepared for qPCR by reverse transcription of 1 μg of RNA using RevertAid H minus First strand cDNA Synthesis kit (Thermo scientific) according to the manufactures standard protocol. A qPCR reaction was prepared using PrimaQuant 2x qPCR-Mastermix (Steinbrenner) by mixing 5 μl of sample (1:10 diluted cDNA) with 5 μl of Mastermix (including 0,3 μM of forward and reverse primer, **Supplementary Table S9**) on a 384-well qPCR plate (LightCycler 480 Multiwell Plate 384, white, Roche). The plate was then centrifuged (2 min, 2000 rpm) and processed for qPCR in a Roche 480 LightCycler using the following PCR program: (i) 10 min at 95 °C, (ii) 15 sec at 95 °C, (iii) 60 sec at 60 °C, repeat step ii) and iii) 40 times and measure fluorescence at 494 nm-521 nm during step iii). Melting curve analysis of each sample was performed to assess reaction quality. Relative expression of each gene in each sample (normalized to rps7 expression) was analysis as log_2_-foldchange over RLUC dsRNA treated samples (Nolan et al., 2006; Schmittgen and Livak, 2008). qPCR primers were designed using the GETprime web service (Gubelmann et al., 2011).

For analysis, all genes in the combinatorial library were annotated manually using FlyBase and literature annotations (Marygold et al., 2013). A more detailed description of all methods including those for supplementary materials can be found in the **Supplementary Methods**.

All code used for the analysis presented in this study is available for download at:

https://github.com/boutroslab/Supplemental-Material/tree/master/Heigwer_2018

All raw data is available at:

https://figshare.com/s/fe82824053056805032e (DOI: 10.6084/m9.figshare.6819557)

## ACKNOWLEDGMENTS

We thank Benedikt Rauscher, Maja Funk, Niklas Rindtorff for critical comments on the manuscript. We thank Luisa Henkel, Tianzuo Zhan and the members of the Boutros laboratory for fruitful discussions and valuable inputs into the study.

